# Promises and pitfalls of *in vivo* evolution to improve phage therapy

**DOI:** 10.1101/816678

**Authors:** James J Bull, Bruce R. Levin, Ian J. Molineux

**Affiliations:** Department of Biological Sciences, University of Idaho, Moscow, ID, USA; Department of Biology, Emory University, Atlanta, GA, USA; Department of Molecular Biosciences, University of Texas, Austin, TX, USA

**Keywords:** mathematical model, evolutionary prediction, dynamics, cocktail, within-host

## Abstract

(Background) Phage therapy is the use of bacterial viruses (phages) to treat bacterial infections. Phages lack the broad host ranges of antibiotics, so individual phages are often used with no prior history of use in treatment. Therapeutic phages are thus often chosen based on limited criteria, sometimes merely an ability to plate on the pathogenic bacterium. It is possible that better treatment outcomes might be obtained from an informed choice of phages. Here we consider whether phages used to treat the bacterial infection in a patient might specifically evolve to improve treatment. Phages recovered from the patient could then serve as a source of improved phages or cocktails for use on subsequent patients. (Methods) With the aid of mathematical and computational models, we explore this possibility for four phage properties expected to promote therapeutic success: *in vivo* growth, phage decay rate, overcoming resistant bacteria, and enzyme activity to degrade protective bacterial layers. (Results) Phage evolution only sometimes works in favor of treatment, and even in those cases, intrinsic phage dynamics in the patient are usually not ideal. An informed use of phages is invariably superior to reliance on within-host evolution and dynamics, although the extent of this benefit varies with the application.

## Introduction

Driven by well-warranted concerns about the growing numbers of infections with antibiotic resistant pathogens, there has been a resurrection of interest in, research on, and even patient trials with a therapy that predates antibiotics by more than fifteen years: bacteriophages [1–4]. Fueling this enterprise, which is now becoming increasingly commercialized (see above references), are well-publicized successes of a handful of compassionate uses of phage to treat chronic, recalcitrant bacterial infections. Patients who were on their way to succumbing to or remaining infected with antibiotic resistant, *Acinetobacter baumannii*, *Mycobacterium abscessus* and *Pseudomonas aeruginosa* survived and in some cases were cleared of the infecting bacteria following treatment with phages [5–8]. The role of phages in these success is not always clear, as these treatments were necessarily uncontrolled and involved single patients. Also not apparent is whether other (and how many) compassionate use trials fail. Indeed, recent clinical trials of phage therapy that did involve controls have often been failures [e.g., 9–11].

Could the difference between success and failure in these therapeutic efforts be simply a matter of phage suitability for treatment – that some phages are better that others? There are certainly properties that *a priori* seem to be ideal candidates for therapy and that might guide choices of phages (Table 1). Against this possibility, host range was the sole criterion for phages used in some patient successes, suggesting that treatment outcomes might be indifferent to phage choice. Yet some experimental work indicates that phage characteristics can influence success [12,13, and see below]. Furthermore, given that therapeutic success with patients often required weeks to months and multiple administrations of phages, it is entirely plausible that a wise choice of phages could vastly improve the progress of treatment, even when the long term outcome is the same for arbitrarily-chosen phages.

**Table 1.**
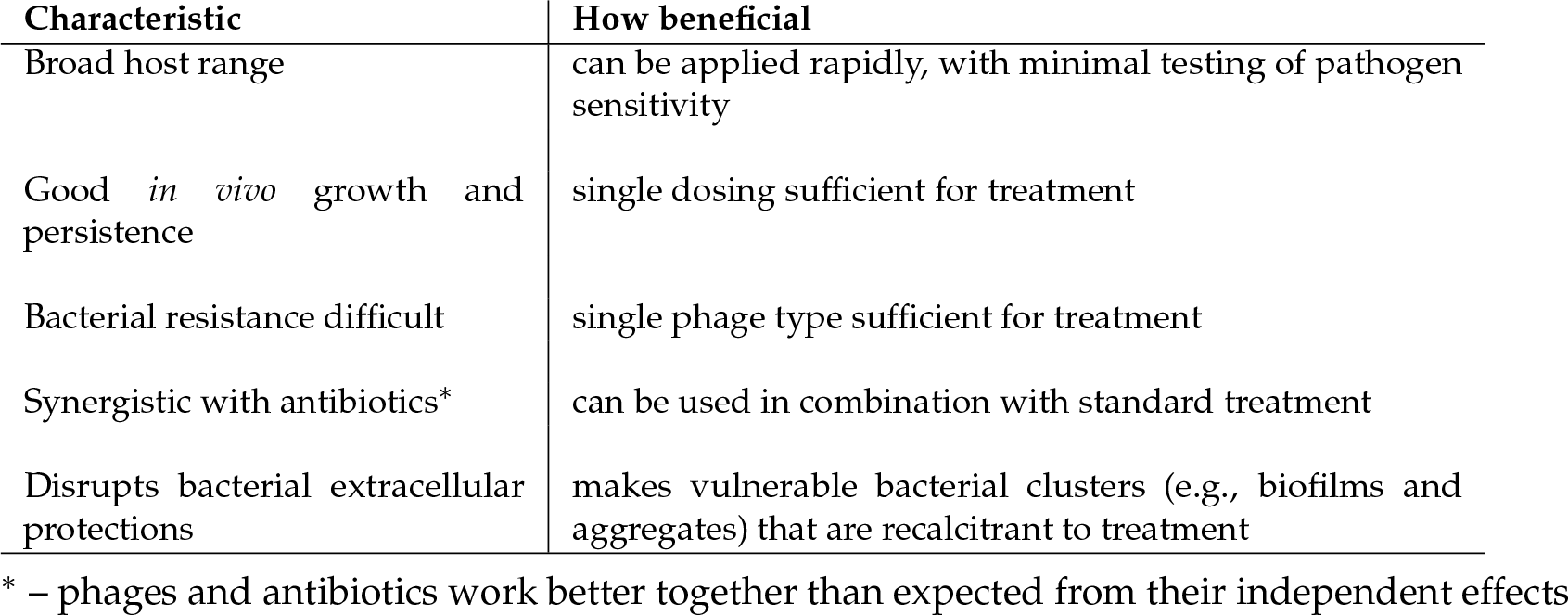
Phage properties ideal for therapy

If phage characteristics matter to treatment success, might there be a shortcut to identifying the best phages? Motivated by the highly visible field of directed evolution of biological molecules [14], it is tempting to consider a directed evolution parallel for phage therapy: can phages evolve themselves to improve treatment? There is even an historical precedent for a similar approach: in his earliest phage therapy experiments, D’Herelle [15] used phages isolated from patients to treat other patients. The idea also has intrinsic appeal: one of the oft-cited benefits of phages over antibiotics is that, by amplifying within the host, they can evolve and perhaps keep abreast of any bacterial resistance evolution. Within-host evolution might then obviate the need to design treatments wisely, instead letting evolution rather than engineers identify good treatment practices. However intrinsically worthy that idea might seem, it is not a simple one to evaluate from first principles: processes of phage dynamics and evolution can be highly unintuitive, and their effects on treatment adds a second layer of complexity. To avoid reliance on intuition, we approach the problem of within-host phage evolution using mathematical and computational models.

Here we critically evaluate the premise that, during the course of therapy, selection favors phages and phage combinations that are specifically effective at killing the bacteria causing an infection. Use of the phage ‘effluent’ from one patient to treat another assumes that the infections in different patients will be suitably similar, and thus possibly of limited utility, but the study of phage evolution from convalescent patients might at least help identify principles that apply to other infections. Given that therapeutic use of phages in humans is so limited at present (at least in the U.S. and much of Europe), initial research on within-host evolution will likely need to use non-human animals and identify principles that could be applied to humans. Indeed, widespread adoption of the One Health perspective, coupled with the problem of drug resistant infections in non-human animals and the need to limit antibiotic use in those settings [16], provides many reasons to develop phage therapy for animals – with the added benefit of helping formulate practices that would work on humans as well. Although established animal infections may prove at least as useful as human infections in discovering principles that work well for treatment, experimental infections may need to be chosen carefully to avoid artefacts.

### A precedent for the need to choose phages wisely

One critical understanding in the development of good phage therapy practices is the extent to which different phages matter to treatment success – beyond the obvious one of whether the phages can grow on the infecting bacterium. If a phage’s ability to plate on a bacterium is all that matters, phage therapy might unfold with nearly the same generality as did antibiotic use. Conversely, if phage characteristics greatly affect outcomes, success may hinge on careful choice of the best phages – unless within-host evolution is self-correcting. Even when different phages are each capable of effecting long term recovery, they might differ in speed of recovery, cost of recovery, and morbidity.

A 1982 study suggested that phage choice can have a profound effect. Using an experimental mouse infection with a K1-capsulated bacterium, Smith and Huggins [12] discovered that lytic phages isolated from the wild fell into two classes regarding treatment success. With simultaneous infection and treatment, one class of phages provided almost 100% recovery, the other closer to 30%. The underlying basis of the difference was that the ‘good’ phages required the bacterial capsule for infection, the ‘poor’ phages did not. The work set a precedent in demonstrating that phage therapy could work *in vivo* and in exploring the dynamic basis of success (although it is less often noted that treatment success fell off dramatically with an 8 hr delay in treatment [12], the cells rapidly becoming recalcitrant to treatment [17]). The work Smith and Huggins [12] was especially important in revealing that differences among phages could profoundly affect treatment success. It thus follows that, if we can identify or even generate good phages, better treatment should follow.

### Phage properties subject to selection in patients

Smith and Huggins [12] sets the stage for our study, but the emphasis here is one step beyond: to what extent can *in vivo* evolution and cocktail competition be used to improve treatment or at least identify the best phages? It is obvious from the Smith and Huggins experiment that the ‘poor’ phages were not themselves evolving fast enough *in vivo* to achieve the high success rates of the good phages, or the rescue rates would not have differed between the phages. But we may instead ask about the effect of within-host phage competition: if both types of phages were injected together into the mouse, would the ‘better’ phage come to dominate the phage population? One can do a formal within-host competition experiment to answer the question for these phages [e.g., 18], but we seek principles that will allow us to predict the outcome more broadly.

At a general level, predicting the outcome of phage competition requires understanding how the entire suite of characteristics for each phage affects its growth and killing of bacteria – i.e., the phage as a whole. It is also possible to consider the evolution of individual phage characteristics that we expect to influence treatment success: (a) phage decay rates [19], (b) enzymatic digestion of a protective bacterial extracellular matrix [20,21], and (c) ability to block bacterial resistance evolution [4]. Considering the phage as a whole might seem to make the most sense, but it may be a single characteristic that determines treatment success, as with the Smith and Huggins [12] example.

A technical note is that we apply the concept of within-host ‘evolution’ in both a narrow sense and a broad one. The narrow sense is the standard Darwinian process of natural selection: mutations arise and ascend based on their merits, eventually resulting in most of the phage population carrying the mutations, all during the interval of treatment. Our broad-sense use is a competition of different phage types within cocktails: the different phage types increase or decrease as a proportion of the total within-host phage pool, their relative fitnesses depending on their intrinsic properties. (For an overview of issues with cocktail pharmacology, see [22].) Both processes will occur in any host treated with a cocktail. The main differences between evolution in the narrow sense versus cocktail dynamics are: (i) with cocktail dynamics, the phages may differ in several characteristics that affect their competition with other phages in the mix, so the selective advantage of one trait may be overwhelmed by other differences, and (ii) because cocktails start with high levels of variation, cocktail dynamics will typically be much faster than Darwinian dynamics. However, reliance on cocktail dynamics to inform and improve treatment can only be applied when different phages are available, but this may not always be the case.

The focus here is two-fold. First, does within-host phage evolution work in favor of treatment? Second, can it work fast enough within one patient to plausibly augment that patient’s outcome? Regardless of the second answer, if within-host dynamics and evolution at least work in favor of treatment, then we might collect phage from a treated patient to use in treating subsequent patients — D’Herelle’s method. Any understanding of these processes is most accurately studied *in vivo*, but *in vitro* work and modeling is often all that is available. In any case, modeling is often necessary to interpret the *in vivo* work. New results provided here are limited to modeling, using a mix of mathematical models and computater simulations. Some processes can be analyzed as simple optimization problems, but others must be embedded in non-linear dynamics that involve interactions and feedbacks. Even the simplest evolutionary problems require quantitation to appreciate the impact on treatment success. Treatment outcomes may be qualitative (infection clearance or not), but the difference between controlling an infection versus bacterial escape may rest on minor quantitative differences, thus requiring a models framework.

## Methods: models

### The standard model and anomalies from phage therapy results

For over half a century, the standard models of phage-bacterial dynamics assumed mass action – full mixing – with homogeneity of bacterial and phage states [23–25]. This model was appropriate for bacterial growth in flasks or chemostats, which was the experimental norm and allowed for easy parameter estimation. The tight coupling between models and experiments and the ease of analysis afforded by this model resulted in most of our current understanding about phages being based on a mass action perspective. The typical outcome is that phages quickly and profoundly depress high densities of sensitive bacteria. From this framework, the attraction of using phages to control bacterial populations is easily understood: a single application of even a small number of phages can virtually wipe out the bacterial population in hours, leaving only a handful of survivors for the immune system to clear.

Full mixing is a poor approximation to phage dynamics in bacterial biofilms and other structured environments. Indeed, in early experiments sensitive bacteria were discovered to have found refuge from phage predation in the walls of chemostats. Aggregates, abscesses and other highly structured populations are now also recognized as important features of bacterial infections [2,26,27]. Furthermore, recent successes with phages in treating individual patients has revealed that some key mass action outcomes are violated [reviewed in 2,4]. First, success in clearing or even suppressing an infection with phages is gradual and sometimes requires months – violating the principle that phages quickly outnumber and kill their sensitive prey. Second, single infusions of phages are often not sufficient: multiple infusions of high doses of phages are required. Both outcomes are difficult to reconcile with the standard model, and although there could be varied causes, spatial structure of bacterial populations (and the associated bacterial inhomogeneity in susceptibility to phages) is one obvious and empirically-justified alternative to consider.

### A dynamics model to accommodate spatial structure

The observations from successful therapies pose a dilemma: high densities of genetically sensitive bacteria persist in the presence of phage. Resolution of this anomaly would seem to require that much of the bacterial population is protected from attack, whether by a polysaccharide matrix, ionic gradients, low receptor expression, or even by some property of the host. Taking as inspiration the high-resolution imaging study of Darch <u>et al.</u> [28] on *Pseudomonas* in a structured environment, we model the bacterial population as consisting of a mix of individual cells and multi-cellular aggregates. The individual cells are fully sensitive to phage attack and obey standard mass action dynamics (and for convenience will be referred to as ‘planktonic’), whereas aggregates are protected from phage attack. However, aggregates convert into planktonic cells and vice versa; this switching of states accommodates the observation that aggregate numbers are somewhat reduced by phage attack but much less so than isolated cells [28]. Our formulation constitutes a refuge model [also true of 26, but we present a more explicit refuge].

Equations and parameters given in Appendix A. For the model of a single bacterial strain (Appendix A1), the main change from the standard model is to allow bacteria to move between a sensitive state of individual cells (given by *B*) and a protected state of aggregates (*B_A_*). Mass action applies to phage-bacterial interactions in the sensitive population, where they are treated as if planktonic. For the model that incorporates resistant bacteria (Appendix A2), each bacterial strain is modeled as if switching between an individual state (planktonic) and aggregates, but all states of the resistant strain are protected against phage infection.

Numerical trials of the equations were run in Mathematica 12.0.0.0, also used to generate the figures. Mathematica files are uploaded as supplements.

## Results

### Growth on a single bacterium

A basic question is how phage evolution works to suppress bacterial numbers. For example, can phage evolution ever allow bacterial density to increase? From a therapeutic perspective, we suppose that decreasing bacterial numbers improves treatment success. In the simplest system – well mixed with a single, sensitive bacterial state – the answer from modeling efforts is straightforward: bacterial evolution may lead to higher bacterial densities, but phage evolution does not. At dynamical equilibrium, one phage type will prevail, the phage that most depresses bacterial density [25]. Dynamic equilibrium is not necessarily applicable to therapeutic success, but a similar theoretical result applies to phages invading a bacterial population: the phage with fastest growth will prevail while bacteria are abundant, which means that selection is for the fastest growth and killing [17]. These results apply equally to phage evolution in the narrow sense and to dynamics of different phages within the therapeutic regime.

A simple use of this principle in a cocktail setting would be ‘phage sorting’ – separating poorly-growing phages from those that grow well. *In vitro* growth – easily determined in advance of treatment – may poorly reflect *in vivo* growth [13], so the appropriate environment is within the patient. Adding a cocktail of phages to an infected patient and then sampling hours or days later should easily determine which phages grow (and survive) at least moderately well in the host. Sorting could be used in a highly quantitative manner, but such refinement is not advisable, as considered below. Phage sorting overlaps with D‘Herelle’s approach of isolating therapeutic phages from convalescing patients, the difference being that here, the phages are being administered to the patient and then collected later. D’Herelle’s method recovered phages that the patient acquired naturally.

The simple nature of phage selection and evolution changes when multiple bacterial states exist – bacterial heterogeneity. We assume that all states are sensitive to the phage and merely differ in phage-infection properties — burst size, adsorption rate, or lysis time. Different bacterial states could be driven by environmental or physiological heterogeneity, as might be represented by planktonic versus aggregate/biofilm/abscess cells, growth on different substrates, or variation in surface molecules, e.g., capsules or O antigens affecting adsorption. These kinds of heterogeneity would develop prior to phage administration and typically have a non-genetic or ‘phenotypic’ basis. (Genetic bacterial resistance evolution is addressed in a different section.)

Well-mixed systems with heterogeneous cell states present several unintuitive outcomes that violate a direct link between phage evolution and treatment success. Notably, selection can favor phages to avoid some types of hosts, even when the phage can productively infect those hosts [29,30]. However, we conjecture that the types of phenotypic variation most relevant to infections will be spatially structured, such as planktonic cells dominating liquid tissues, and biofilms, aggregates or abscesses in solid tissues. With spatial structure, different phage types or mutants will differentially amplify in patches of cells where they grow best. In these settings, phage evolution and competition should often work loosely in favor of treatment, at least on a local level, but dynamical complexities allow for exceptions.

With phage cocktails or *in vivo* evolution of phage mutants, the composition of actively growing phages exuded from patients will typically change over time as bacterial densities are suppressed at different rates in different locations. The cumulative phage composition should mirror the phages with the biggest numerical effect on bacterial killing at the time. However, one dilemma faced in phage collection from the patient is that the relative abundance of different phages in the collection need not match phage importance to reduction of the patient’s symptoms. If the bacteria responsible for morbidity or maintenance of the infection are confined to small or semi-protected bacterial populations, the patient’s phage output may be dominated by phages that kill large numbers of bacteria in other tissues that contribute only slightly to virulence. Phage evolution itself will be driven by growth on the larger populations of bacteria in the host. As a consequence of these varied processes and possibilities, there is no predicted time to harvest phages from one patient that would ensure maximum benefit for use on subsequent patients.

#### Conclusions

Intrinsic processes of phage evolution work unambiguously in favor of treatment when bacteria exist in a single, well-mixed state. When multiple bacterial phenotypic states exist and are well mixed, phage evolution and cocktail competition can give rise to unintuitive outcomes whereby phages are selected to avoid some bacteria. Spatial structure of bacterial states, as with biofilms (aggregates or abscesses) versus planktonic cells, may align phage evolution with suppressing bacterial numbers locally, but these processes are not easy to study and are not well understood *in vivo*.

Although encouraging, these results do little more than suggest an expected, qualitative direction of evolution regarding treatment. They offer nothing on the speed or magnitude of infection clearance. Nor do we have any sense of how much within-host evolution to expect, given a potentially short duration of treatment. Deeper insight to the uses of *in vivo* evolution to improve treatment comes by addressing specific phage characteristics, next.

### Phage decay rates

Using two well-characterized tailed phages (*λ* and P22), Merril <u>et al.</u> [19] showed that (i) the initial or ‘wildtype’ isolate of each phage was rapidly cleared from mice (in the absence of bacteria), and (ii) mutants of each could be isolated that were cleared more slowly (the clearance rate evolved to approximately 20% of the initial). The study further showed that the long-persisting phages were advantageous in prophylactic therapy – administered in advance of the infection – presumably from their ability to persist up to the time of bacterial introduction. The interval in which phage decay was measured was short, less than a day after inoculation, too fast for adaptive immunity to develop in the naive mice. The suggestion was that phage clearance was due to the reticuloendothelial system.

These results raise two questions. (i) Is the benefit of long-persisting phages large enough to affect treatment outcomes? If phages amplify during treatment, phage differences in survival might have little benefit, except when phage amplification is weak. (ii) Will long-persisting phages evolve during treatment and evolve quickly enough that there is little to be gained by inoculating long-persisting phages?

Of the phage characteristics analyzed in this paper, this one experiences the simplest evolutionary process: each phage type and mutant evolves independently of other phages, and the selection does not involve frequency- or density-dependence. Simple calculations can be informative as a first step. Let phage survival follow exponential decay, e^−wt^ for the ‘wildtype’ and e^−mt^ for the mutant (*m* < *w*, with *t* in minutes). If the mutant starts at frequency *p*_0_, the ratio of the mutant to wildtype will change in time according to *R*(*t*):

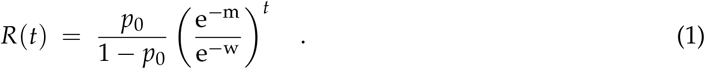

R(t) increases without bound over time, but only as long as both phage types are abundant enough to approximate deterministic dynamics.

The time at which the mutant comprises half the phage population is approximately

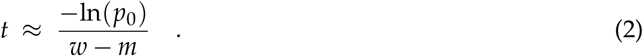

For new mutations, the numerator will typically lie in the range of 10-20. Although this calculation ignores phage amplification (which should have little impact on *relative* phage abundances in the absence of other differences between the phages), this formula shows that the time for an initially rare mutant to dominate the phage population depends on the reciprocal of its relative advantage (*w* − *m*). Given that both rates must be positive (and *w* > *m*), the magnitude of *w* − *m* cannot exceed *w*. A wild-type with a very low clearance rate (*w* ≪ 1), cannot be quickly displaced even for a mutant that is never cleared (*m* = 0). Evolution works fastest the more important the change, but of course, if *w* is small, there is little benefit from further reductions.

Are clearance rates of wild phages high enough to evolve during an infection? Calculations from the results in Merril <u>et al.</u> [19] and Westwater <u>et al.</u> [31] gave wildtype clearance rates on the order of 2 − 6 × 10^−3^ [see calculations in 32]. A 5-fold improvement (say from 5 × 10^−3^ to 1 × 10^−3^) and *p*_0_ = 10^−6^ requires just over 2 days for the mutant to comprise half the population. This may be a long time to wait for any treatment benefit afforded by the mutant. Furthermore, if the phage population was declining (e.g., unable to maintain itself), the phages might be too rare after 2 days to observe any improvement in treatment from mutant ascendance. An interesting observation is that phages engineered by phage display (engineered to carry exposed, short peptides on the outside of the virion) may be particularly prone to rapid clearance [33], as if there is a strong selection to eliminate the engineering.

When phages are used prophylactically, it is obvious that any benefit of reduced clearance must be achieved by evolving the phage before administration. The goal is to maintain the phages as long as possible in a non-growing state in anticipation of a possible bacterial infection. The drawback of starting with a rapidly-decaying phage cannot be overcome by mutation and phage evolution when the phage are not growing. However, the situation changes if phages are administered to an infection, because now a mutant, slow-decaying phage may arise and could in principle take over the within-host population rapidly. Were this the case, one could rely on within-host evolution to improve treatment. To address this possibility, we rely on a numerical model (Fig. 1). As described in our Methods section, this model attempts to capture more realistic dynamics than is typical of the standard ‘mass action’ model of phage-bacterial dynamics. This new model assumes both a sensitive bacterial population and a ‘refuge’ population of bacteria protected from infection, leading to reduced oscillations and (sometimes) a need to serially inoculate repeated doses of phage to keep bacterial numbers low. This new model is inspired by observations from phage therapy patients that phages may be unable to maintain themselves [2, and see Methods]. With this model, it is seen that (i) decay rate matters to rapid bacterial suppression, and (ii) early in treatment, evolution is not as effective as is dosing with the pure, slowly-decaying phage (Fig. 1).

**Figure 1.**
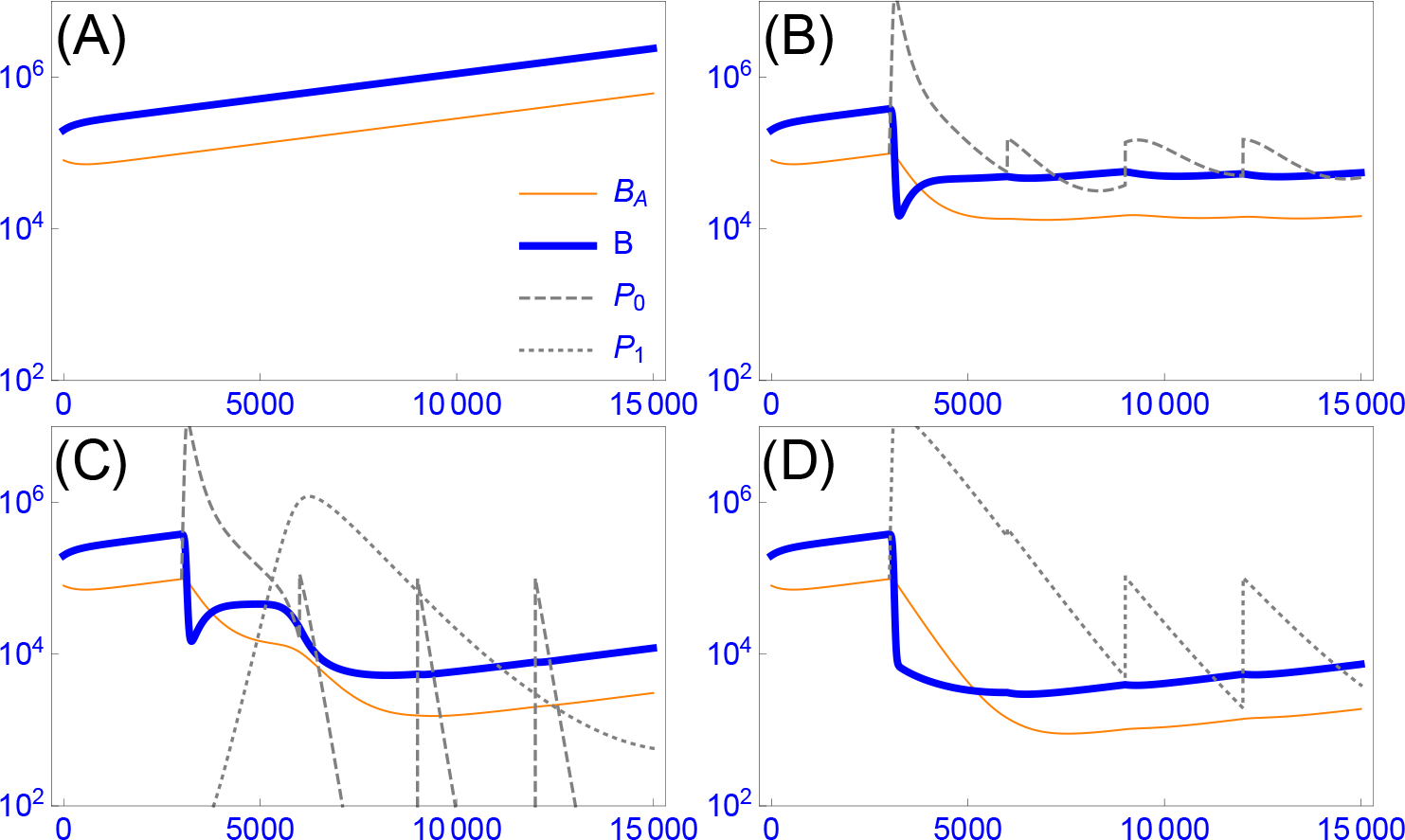
Phage dynamics and evolution as a property of phage decay rate. The vertical axis gives density, the horizontal axis time (minutes). Thin orange curves are protected bacteria in refuges, thick blue curves are susceptible, planktonic bacteria. Dashed grey indicates the fast-decaying phage (*P*_0_, decay rate of 0.008 per min), dotted grey indicates the better, slow-decaying phage (*P*_1_, decay rate of 0.002 per min). The inset key (A) applies to all panels. All trials use the same bacterial growth parameters and initial bacterial densities (given in Appendix A1). When phage are present, they are first added at 3000 minutes and added every 3000 minutes thereafter. (A): Bacterial densities slowly increase in the absence of phages. (B): Treatment with 10^5^ rapidly-decaying phages causes a sudden decline in free bacterial densities, a somewhat slower decline in aggregate bacteria. The system is approaching equilibrium in that phage and bacterial densities are being maintained. (C): The phage inoculum consists of 10^5^ rapidly-decaying phage and 0.1 slowly-decaying phage, the latter value to represent mutation. The slowly-decaying phage ascends profoundly but then drops when bacterial densities are too low to sustain it. (D): the phage inoculum consists of just 10^5^ slowly-decaying phage. There is a substantial difference between the treatments with a pure fast-decaying phage or a pure slow-decaying phage. The main effect of starting with a slow-decay phage instead of relying on within-host evolution lies in the early dynamics, although a modest lingering benefit is apparent. Parameter values and initial conditions are given in Appendix A1. Outcomes vary with parameter values, and the actual effect of within-host evolution or treatment with pre-evolved, slowly decaying phages would need to be evaluated for each specific application.

#### Conclusions

If the wildtype phage has a suitably high decay rate, evolving the phage to a lower decay rate in advance can offer a significant improvement in treatment and is the only solution to improve phage prophylaxis. The treatment benefit of such pre-emptive evolution depends on the wildtype decay rate and the magnitude of improvement afforded by available mutations. Relying on within-host evolution to achieve dominance by a slow-decaying phage becomes feasible only when the treatment phages actively grow and maintain themselves in the host for days. However, phages recovered from a patient, as in D‘Herelle’s approach, may well have evolved lower decay rates and could be used with subsequent patients.

### Matrix degrading activities: depolymerases

Some phages encode enzymes (depolymerases and lysins) that may have important direct or indirect antibacterial activities. Depolymerases (the focus here) degrade bacterial secretions and may foster bacterial killing and clearance [20,21,34–38]. Indeed, it is now known that the superior phage type discovered by Smith and Huggins [12] differs from the poor phage type by encoding a depolymerase that degrades the bacterial capsule [39]. The free enzyme is likely critical to treatment success – the pure depolymerase can successfully treat an infection even in the absence of phage [40–44]. Protective bacterial secretions include more than just bacterial capsules, such as the chemically complex matrices in biofilms and aggregates [21,28,37,45–47]. Phage-encoded enzymes do not necessarily exist for all bacterial matrix components, but enzyme-encoding genes from other sources (bacteria, fungi) can be engineered into and expressed from phages to achieve degradation [e.g., 21,48,49].

In wild phages, depolymerases are assembled as part of the phage particle (virion): the tailspikes used in the initial, specific recognition of the host. The enzymes allow the phage to penetrate the protective surface layers of the bacterium and thus provide a direct benefit to the individual phage much as does any phage-encoded contribution to infection. But depolymerases also provide a dispersed benefit for treatment: considerable free enzyme, that not assembled onto virions before cell lysis, is released and can diffuse into the surrounding environment where it acts separately from the phage: the effect of the diffusing enzyme can often be observed on plates as an expanding halo beyond the plaque [45,50–52].

Interest here is primarily in the dispersed benefit of phage-encoded enzymes – a benefit beyond the phage killing itself. Free enzyme augments other forms of bacterial killing by degrading the protective layers around many bacteria. Loss of those layers exposes the bacteria to attack by complement, immune cells, drugs, and even by other types of phages that would not otherwise be able to access the bacteria. From a treatment perspective, free enzyme is ideal because it augments treatment by multiple antibacterial agents and immunity. Furthermore, genetic engineering greatly expands the capacity of phages to degrade substrates that do not directly aid infection [21]. If this property of a phage or cocktail can be evolved and, importantly, be maintained, it has the potential to greatly augment treatment.

Evolution of a dispersed effect from an enzyme is not straightforward. Released enzyme diffuses into the local environment and acts as a ‘public good’ that can benefit phages that do not produce the enzyme. By benefiting others, the producer phage is subjected to a ‘tragedy of the commons’ that works against its success in competition with non-producing mutants and phage types [52,53]. The ‘tragedy’ can be especially acute for phages engineered to encode depolymerases, as those enzymes are usually encoded entirely as free enzymes and not part of the virion [21]: evolution can quickly dispense with the engineered gene [53]. Enzymes that are encoded as part of the virion are far more prone to be genetically (evolutionarily) stable, but the benefit the excess enzyme provides other phages nonetheless works against their success in a cocktail. Maintenance of an enzyme-producing phage in cocktails has little to do with its public-good benefit in clearing the infection, and such phages can be lost even if bacterial clearance depends on their presence [52]. Evolution and dynamics can work in favor of enzyme-producing phages when the bacterial environment is spatially structured, where the free enzyme may not diffuse far to help other phages [53], but there is yet no evidence on whether appropriate spatial structure applies to natural infections.

#### Conclusions

Although the dispersed benefits of enzyme-producing phages can greatly enhance clearance of structured bacterial populations, within-host evolution and competition among phages are not aligned with maintaining enzymes for that reason. Phage competition will not typically improve the collective enzyme-degrading activities of the input phages. Maximizing the treatment benefit from engineered depolymerases requires that the engineered phages also be designed to avoid evolutionary loss of the enzyme. Cocktails should be administered with an *a priori* understanding of dynamics that may work against them.

### Phage evolution to overcome bacterial resistance

The most-studied evolutionary process with phages is their impact on and response to bacterial resistance evolution. The problem has long been considered from the perspective of genetic and molecular mechanisms [23,54–58] and as phenotypic outcomes of phage-bacterial arms races in co-culture. Arms races have been explored both in conditions of natural or semi-natural dynamics [59–64] as well as highly contrived conditions in which vast numbers of phage were amplified on a permissive host and forced to grow on resistant hosts [54–57]. Our interest here is in observations from (semi-)natural dynamics.

The ascent of resistant bacteria within the patient will typically be inimical to treatment success and should be avoided. Furthermore, there remains the formal possibility that a phage-resistant bacterium will have increased virulence, perhaps by becoming mucoid and thus less sensitive to immune system components; however, reports of such outcomes so far seem to be lacking. Experimental work indicates that evolution of bacterial resistance is a seemingly ubiquitous and easy response to a high abundance of phages. From *in vitro* co-culture studies of phages grown on single bacterial strains in simple media, a common outcome of arms race evolution is ultimate dominance by resistant bacteria [60]. The arms race may involve a few steps of phages evolving to overcome bacterial resistance and new bacterial resistance evolving, but ultimately a bacterial resistance evolves that cannot be overcome by phage evolution; phage are subsequently lost or their abundance greatly suppressed. This simple story is often violated when bacteria are grown under more complex *in vitro* environments or in natural ones: resistant bacteria may fail to ascend, with the phage persisting and permanently suppressing the bacteria [65,66]. However, other outcomes have also been observed [61,62,64]. The hope is for *in vivo* outcomes that avoid bacterial resistance, but in fact, resistant bacteria have been observed to ascend within patients [67]). So the hope then turns to learning how to choose phages to direct the outcome toward blocking bacterial resistance evolution.

### Intrinsic phage evolution to overcome bacterial resistance is not assured

Bacteria are selected to avoid killing by phages. Likewise, phages are selected to overcome bacterial resistance, both *in vivo* and *in vitro*. Nonetheless, reliance on *de novo* mutation and Darwinian evolution to overcome bacterial resistance faces two problems. First, the requisite mutations may not arise – the jump to utilize a new bacterial receptor may require an improbable combination of multiple mutations in the phage genome. Second, even when appropriate phage mutations arise, their ascent may rely on resistant bacteria having become abundant enough to rapidly amplify the mutants. The population dynamics work against evolution because of a mismatch between selection and phage population size (opportunity for mutation): selection is strongest when the resistant bacterial population is largest, but the phage population may plummet due both to a reducing number of sensitive hosts and to clearance before resistant bacteria become common. These dynamics commonly lead experimentalists to evolve new phage host ranges by alternately allowing amplification on permissive hosts [e.g. 54,55].

### Cocktails can be designed to block stepwise bacterial escape, but they can experience lags and phage loss

Natural phage evolution is not the only solution to bacterial resistance. One alternative is to isolate wild phages that grow on the resistant hosts, then use those in the therapeutic cocktail. Engineering is another possibility for generating resistance-blocking phages [49,68–70]. Advance selection of resistance-blocking phages is possible because bacterial evolution to resist a single phage often follows a common molecular pathway, enabling the investigator to replicate *in vitro* the within-patient arms race. One simply evolves resistant bacteria *in vitro* and then selects resistance-blocking phages in advance of treatment, ultimately including them in the cocktail at sufficiently high concentrations that they are not lost before substantial numbers of resistant bacteria have arisen. Indeed, our mechanistic understanding of resistance evolution is so advanced as to realize that phages infecting the same host by using different receptors will provide complementary blocks to bacterial resistance evolution — pathways of resistance evolution can thus be anticipated even before the patient experiences the resistant bacteria. Unfortunately, the approach of designing a cocktail to anticipate bacterial evolution may only be applicable to chronic infections, as the time and effort to identify phages and their specific receptors, and then formulating an appropriate cocktail, is considerable.

The dynamics of multiple, resistance-complementing phages in a cocktail works in favor of suppressing bacterial resistance, but the timing is not ideal. The main complication is that phage dynamics are not anticipatory, responding to what is present rather than what is about to occur. Given the intrinsic differences in growth likely to exist among wild phages, the initial dynamics of the cocktail will typically be dominated by one phage, with the other phages dropping to low numbers (or even disappearing) before resistant bacteria ascend [Fig.2B and 71]. Resistant bacteria will eventually ascend and potentially exacerbate symptoms before being suppressed by other phages in the cocktail (Fig.2C), and the cycle may repeat. If the cocktail contains resistance-blocking phages, it would seem that the ascent of resistant bacteria could be avoided by periodic dosing with the cocktail, but a very high dose may be needed (Fig. 2, compare D and E). Overall, therefore, cocktail dynamics can ultimately work in favor of overcoming bacterial resistance (provided that appropriate phages are present), but the best treatment may require an informed design and application of cocktails.

**Figure 2.**
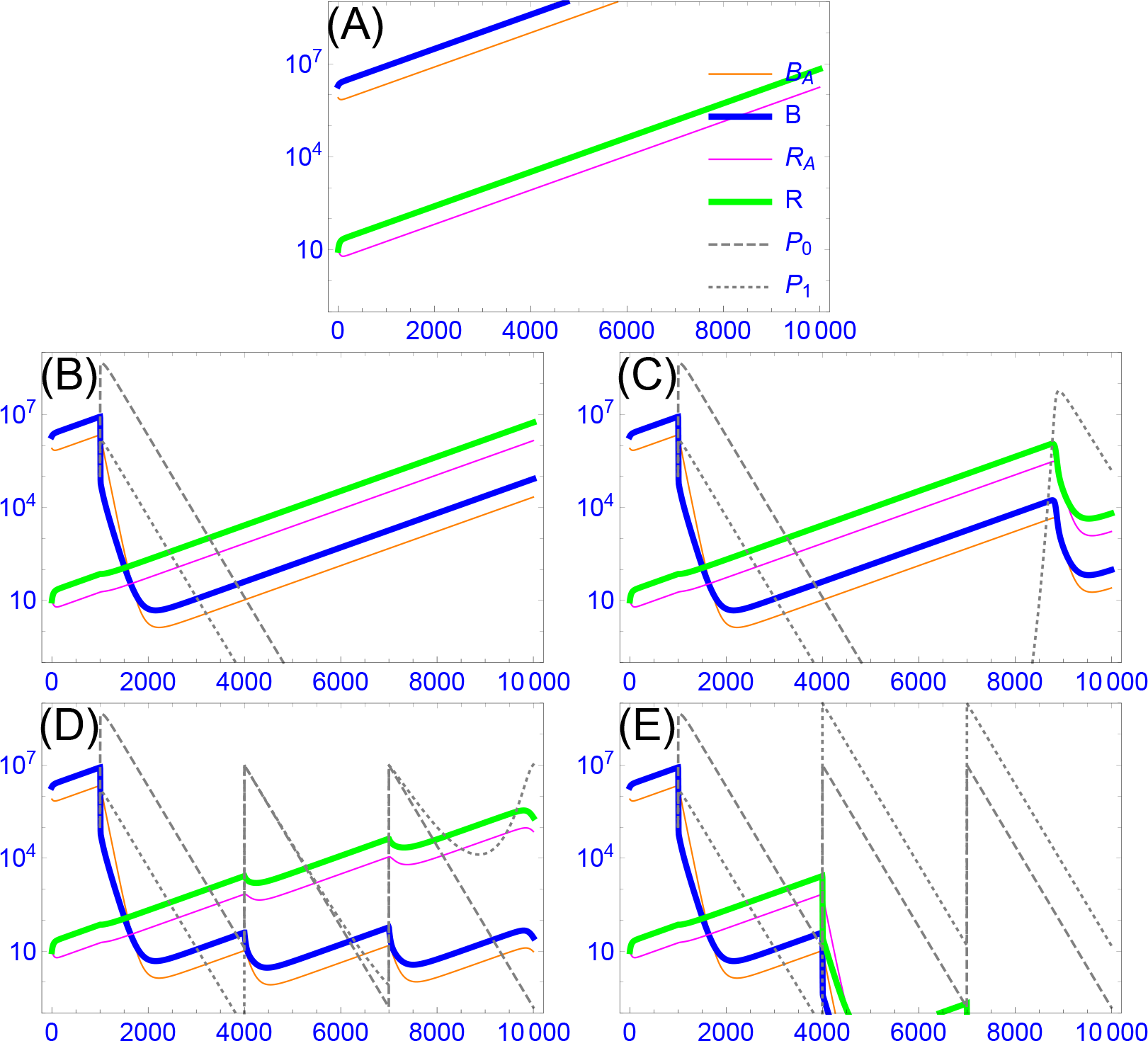
A ‘broad’ host-range phage that blocks bacterial resistance is subject to delayed ascendency. Periodic dosing is only moderately more useful in suppressing bacterial resistance than is relying on intrinsic dynamics. The model assumes two strains of bacteria, each in two states (solid colored curves) and two strains of phage (dashed gray); the inset legend in (A) applies throughout, with equations and parameters given in Appendix A2. The vertical axis gives density, the horizontal axis time (minutes). The phages differ in whether they can infect both bacteria (the ‘broad’ phage, given by *P*_1_) or just one bacterium (the ‘narrow’ phage, given by *P*_0_); the narrow phage has the advantage of a slightly higher adsorption rate. The bacterial strain given by blue and orange curves is sensitive to both phages, the other (green and pink curves) is resistant to the narrow phage. Each bacterium exists both planktonically (thick curves) and in aggregates (thin curves), with aggregates being protected from all phages. The two bacteria differ only in sensitivity to the phages. (A): Growth of the bacteria in the absence of phages. (B): Both phages are introduced at time t=1000 at a density of 10^5^ but are considered to be extinct when densities drop below 0.1. They have a rapid effect of driving the sensitive bacterial strain to low numbers, allowing the resistant bacterium to become the majority. Both phages are lost when the bacterial density is too low to sustain them, and all bacteria begin to recover, maintaining their relative abundances. (C): The same as in (B), e1xcept that phages are never considered to be extinct. The broad phage eventually rebounds in response to the high numbers of ‘resistant’ bacteria, and it suppresses both strains. (D): A cocktail of both phages is applied, each at a dose of 10^5^ (time 1000) and then each at a dose of 10^7^ (times 4000 and 7000). The narrow-host range phage gains early because of its superior adsorption rate. Resistant bacteria eventually ascend and allow the broad phage to maintain itself. Note that there is a substantial lag before the broad phage dominates. (E) The same as in (D), but the inocula at times 4000 and 7000 are increased to 10^9^ of the broad phage. All bacteria are now pushed to near extinction.

The successive collection of phages from the patient should reveal much about the dynamics before and during bacterial resistance evolution. Recovered phages may be a source of improved phages that can be used on other patients infected with the same strain, as may be likely in an epidemic. Such phages may also have been selected for lower rates of clearance by the reticuloendothelial or other systems.

### Resistance-proof phages can avoid evolutionary arms races, but *in vivo* dynamics do not ensure their ascendancy in cocktails

Resistant bacteria do not invariably ascend [4,12, and references above]. Failure to ascend occurs when bacterial resistance seriously impairs bacterial growth in the local environment. Thus, Smith and Huggins [12] observed that their ‘good’ phages required the bacterial capsule for growth; bacterial resistance resulted in loss of the capsule, which rendered the bacterium susceptible to the immune system and thereby prevented ascendancy. Other trade-offs between bacterial resistance to phages and bacterial growth are known; they often depend on the environmental context [4,72]. These observations suggest that single phages might be chosen wisely to prevent the ascendance of resistant bacteria. Use of such ‘resistance-proof’phages has an obvious advantage, even over cocktails, as such treatments do not face the even temporary ascendance of resistant bacteria (although resistance-proof phages are subject to the same predator-prey cycles of other phages). A resistance-proof strategy has even been employed successfully with patients [4,8]. As a possible source of resistance-proof phages, narrow host-range phages may be evolved to overcome bacterial resistance. This selection works well in a laboratory setting where defined cultures are employed, and may lead to resistance-proofing because most host range mutations confer an expansion of host range, rather than a switch. However, the result is usually a compromise, with the rate of adsorption to both the old and new hosts being less than specialist phages that only infect one or the other strain. Reduced adsorption may allow sensitive bacteria to persist at moderate densities or in refuges that might be suppressed by other phages; reduced adsorption will generally affect therapeutic success if rapid phage multiplication is required *in vivo*.

Although resistance-proof phages have sometimes been chosen *a priori* by design [4], a scientific basis for choosing them may not always be available in advance of treatment. Might *in vivo* competitions be used to dynamically evolve such phages? That is, if a cocktail of phages is used in treatment and the cocktail were to contain a resistance-proof phage, would it automatically ascend and displace the others? Not necessarily, at least not because of its resistance-proof status. At the outset, a resistance-proof phage has no advantage when the cocktail is first introduced on the sensitive strain, and whichever phage grows the fastest will predominate [e.g., 71]. When the bacterial population evolves resistance to this first phage – a potentially slow process if bacteria exist in protected states (e.g., Fig. 2) – the resistance-proof phage may be one of several remaining phages capable of growing on the resistant bacteria. Again, the fastest-growing phage will predominate, not necessarily the one that is resistance-proof. Beyond this phase, there are countless possibilities from phage competitions, bacterial competitions, bacterial protected states, and delayed, density-dependent dynamics to preclude generalities. We thus lack assurance that a resistance-proof phage will prevail. Treatment with a single phage known to be resistance-proof appears to be the surest way to maintain a lasting block against bacterial resistance evolution, but the phage must be chosen intelligently rather than by relying on within-host evolution.

Our presentation of bacterial resistance has implicitly emphasized changes in surface receptors affecting phage adsorption. Other mechanisms of bacterial resistance to phages are known [58]. Not all are prone to evolve *in vivo* (e.g., restriction-modification systems of bacteria will not evolve under short-term selection by phages). Furthermore, changes in surface receptors may evolve in deference to other changes [73]. Nonetheless, the dynamics of phage evolution to overcome bacterial resistance by other mechanisms will face the same types of issues identified here.

#### Conclusions

Within-host evolution favors bacterial resistance to phages, but the ascendance of resistant bacteria then favors phages that grow on the resistant strains. Within-host evolution of single phages is not assured of overcoming bacterial resistance, if only because the requisite mutations may not occur. Cocktails may be designed to anticipate and block bacterial resistance evolution, but timing remains a problem: phage evolution in response to resistant bacteria necessarily lags the ascendance of resistant bacteria. Repeated dosing may be required to prevent the temporary ascent of resistant bacteria. Resistance-proof phages offer a solution to a bacteria-phage arms race, but there is no *in vivo* protocol that ensures automatic selection of single, resistance-proof phages, even when a resistance-proof phage is present in the initial cocktail. They must therefore be chosen from experience or an *a priori* understanding of the costs and benefits of bacterial resistance to phages.

## Discussion

It is already clear that, whatever its ultimate utility, phage therapy of bacterial infections will not afford the simplicity of treatment that has been true of antibiotics. Perhaps phage treatments will need to be tailored to the infecting strain and may even need to be tailored to the nature of the infection. However, one hope for phage therapy that does not apply to antibiotics is evolution. When starting a treatment with one or more phages that infects the pathogen, either Darwinian phage evolution or cocktail dynamics within the patient might be aligned with treatment success and ‘automatically’ yield the best phages, obviating the need for a sophisticated understanding of best treatment practices.

Evaluating the potential utility of within-host evolution has special merit in light of the fact that current treatment protocols have been rather simplistic. This is not a criticism, one needs to know what the problems are before designing more complex protocols. In some cases, phages were selected for treatment based on no more than a demonstrated host range *in vitro*. In other cases, phages were chosen merely to grow on the infecting strain and also block the ascent of resistant bacteria, again based on *in vitro* assays. Although infections have been successfully suppressed by some of these protocols, improvements may be warranted to shorten recovery periods or to depress bacterial densities beyond that which was attained. Furthermore, *in vivo* phage evolution may not only augment treatment, but studying it may lead us to new insights that could improve treatment.

The thesis that *in vivo* phage evolution is aligned with treatment success applies to some phage properties but not others. The reason for the different effects of within-host evolution on phage improvement is that the different phage properties evolve under different dynamic processes, and only some processes are aligned with treatment. Properties improved by within-host evolution are phage growth rate and (reduced) clearance rate, as well as the ability to overcome bacterial resistance. However, phage evolution to overcome bacterial resistance has a time lag problem: phage evolution tracks bacterial resistance, so resistant bacteria ascend before being suppressed again. Resistance-proof phages avoid the emergence of resistant bacteria if they are the only phages administered, but when given as one of several phages, they do not ensure against the cyclical rise of resistant bacteria. Wise cocktail design and delivery timing is the more assured way of limiting the ascent of resistant bacteria.

Intrinsic dynamics/evolution within the patient offer no assurance for the maintenance of phage-encoded enzymes that degrade extra-bacterial substances. The release of free, phage-encoded enzymes may profoundly help clear and infection, but this benefit does not help maintain the phages producing the enzymes – unless the infection is highly structured. Again, wise choice of phages, cocktail composition, and dosing times may be required to optimize treatment.

Phages may act in synergy with antibiotics [74–76] and other therapies, although there appears to be no grand generality – some drug-phage combinations are synergistic, others antagonistic [77]. The performance of individual phages may thus well be affected by interactions with other therapies. Indeed, phage therapy with humans may invariably require co-treatment with antibiotics, so phage-drug interactions may well be critical to success. However, we do not expect the alignment of within-host evolution and treatment success to change because of other administrations, so the principles proposed here should not be affected.

### Depressing bacterial densities versus improving infection outcomes

The perspective here has assumed that decreasing bacterial numbers invariably improves the patient’s outcome. Depressing bacterial numbers will no doubt be essential to cure, but there are hints that the relationship between bacterial killing and treatment success is not always straightforward: bacterial lysis releases endotoxins. In an experimental mouse model, lethal but non-lysing phages given at very high doses were shown to improve host survival over lytic phages [78]. Although that experimental system was highly artificial, the result fits with known principles of the immune response, and it points to a wider realm of possible deviations from a simple relationship between bacterial killing by lysis and treatment improvement. For example, treatment with a phage that releases capsular depolymerases may boost immune-mediated killing, which does not entail lysis [e.g., 40,41,43,44].

A second possible concern with ‘successful’ phage therapy is whether bacterial resistance to phages might increase virulence. Bacterial resistance to phages often entails a reduction in bacterial growth rates or densities, but if bacterial density is not the sole determinant of virulence, then an increase in virulence is possible despite reduced bacterial growth. Indeed, some extremely slow-growing bacteria cause serious infections. Bacterial resistance to phages may impart simultaneous protection against immunity, as with bacterial mucoidy. It is thus conceivable that some virulent bacterial mutants may have an advantage within the host only when favored by phage pressure, in which case they would not be observed in the absence of treatment. Although this possibility is speculative, there are unconfirmed reports of symptoms made worse by phage treatment. One conceivable scenario is where the use of broad host range phages and/or complex phage cocktails inadvertently targets benign commensal bacteria that themselves are antagonistic to proliferation of the pathogen.

## Author Contributions

All three authors contributed to all aspects of the paper, except that JJB wrote the Mathematica files to generate the figures.

## Acknowledgments

We thank Steve Abedon for comments and insight to the literature.

## Appendix A: Model details

The standard model of bacterial-phage interactions is one of ordinary differential equations assuming mass action, whereby the number of infections occurring per unit time is simply the product of the phage and bacterial concentrations [scaled by an adsorption rate parameter, 24,25]. This type of model typically leads to a rapid and profound depression of bacteria, followed by a bacterial resurgence and ensuing oscillations of phage and bacterial densities. These dynamics are apparently not representative of phage therapy patients [2]. We instead offer two models of ordinary differential equations that deviate from the standard model and can generate some key outcomes mirroring those of observed with phage therapy: bacteria exist in protected states, phage may slowly decay during treatment, and multiple dosing may be required [2]. Although our models explicitly include many components (minimally 4 variables and 9 parameters), they is proposed heuristically in that they cannot be empirically parameterized. Instead, many parameters need be chosen so that the baseline behavior allows phage to have only a moderate effect on bacterial densities in the short term, and indeed for some trials, that multiple phage inoculations are needed to depress bacterial densities to low levels. Any numerical trial of this model is useful chiefly in illustrating possibilities that are incompatible with the standard model.

## A1. One bacterial strain with two bacterial states and two phages

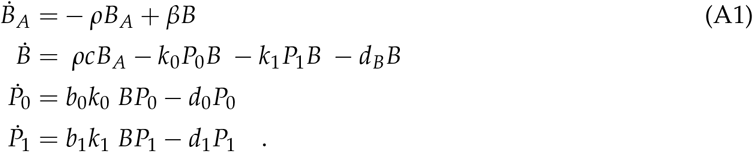

**Table A1.**
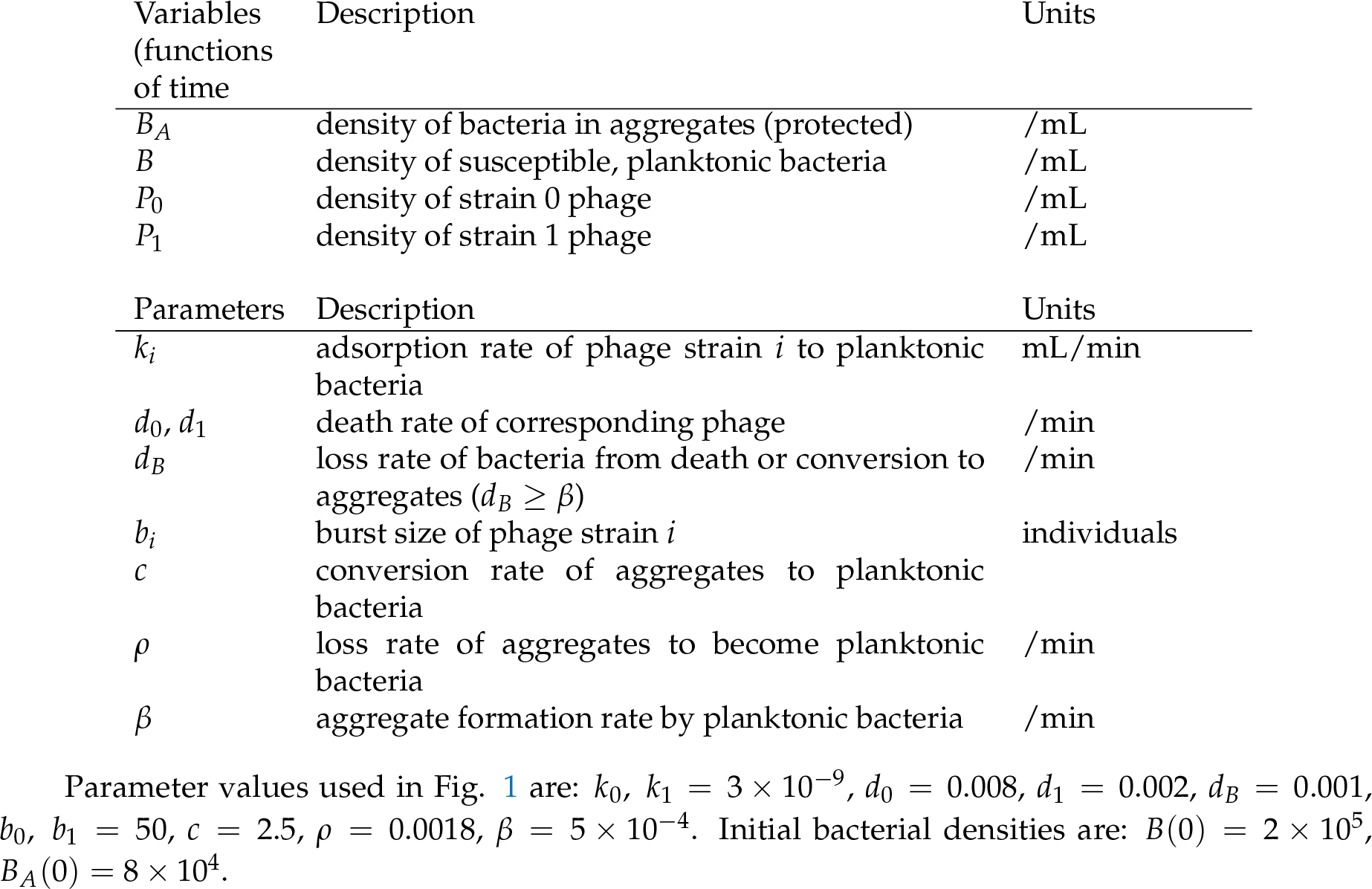
Model variables and parameters

| Variables<br>(functions<br>of time) | Description | Units |
| --- | --- | --- |
| $B_A$ | density of bacteria in aggregates (protected) | /mL |
| $B$ | density of susceptible, planktonic bacteria | /mL |
| $P_0$ | density of strain 0 phage | /mL |
| $P_1$ | density of strain 1 phage | /mL |
| Parameters | Description | Units |
| $k_i$ | adsorption rate of phage strain $i$ to planktonic bacteria | mL/min |
| $d_0, d_1$ | death rate of corresponding phage | /min |
| $d_B$ | loss rate of bacteria from death or conversion to aggregates ($d_B \geq \beta$) | /min |
| $b_i$ | burst size of phage strain $i$ | individuals |
| $c$ | conversion rate of aggregates to planktonic bacteria | |
| $\rho$ | loss rate of aggregates to become planktonic bacteria | /min |
| $\beta$ | aggregate formation rate by planktonic bacteria | /min |

Parameter values used in Fig. 1 are: *k*_0_, *k*_1_ = 3 × 10^−9^, *d*_0_ = 0.008, *d*_1_ = 0.002, *d*_*B*_ = 0.001, *b*_0_, *b*_1_ = 50, *c* = 2.5, *ρ* = 0.0018, *β* = 5 × 10^−4^. Initial bacterial densities are: *B*(0) = 2 × 10^5^, *B_A_*(0) = 8 × 10^4^.

## A2. Two bacterial states with two bacterial states and two phages

This model is similar to that in A1, except that it adds a second strain of bacteria, one that is resistant in the planktonic state, necessarily also resistant as aggregates. To keep the emphasis on the effect of resistance, many of the same parameter values are used for the two types of bacteria and for phage infection of the bacteria.

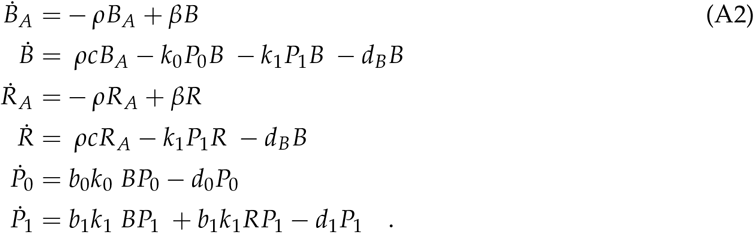

Only variables that differ from the above are defined here. Parameters are the same as above in A1

**Table A2.** New model variables

| Variables<br>(functions of<br>time) | Description | Units |
| --- | --- | --- |
| $R_A$ | density of resistant bacteria in aggregates | /mL |
| $R$ | density of resistant, planktonic bacteria | /mL |

Parameter values used in Fig. 2 are the same as for Fig. 1 except: *k*_1_ = 1 × 10^−9^, *d*_0_ = *d*_1_ = 0.006, *d*_*B*_ = 0.009, *c* = 2.2, *ρ* = 0.018, *β* = 5 × 10^−3^. Initial bacterial densities are: *B*(1) = 2 × 10^6^, *B*_*A*_(1) = 8 × 10^5^, *R*[1] = *R*_*A*_[1] = 10.

© 2019 by the authors. Submitted to <u>Viruses</u> for possible open access publication under the terms and conditions of the Creative Commons Attribution (CC BY) license (http://creativecommons.org/licenses/by/4.0/).

## References

1. Brüssow, H. Phage therapy for the treatment of human intestinal bacterial infections: soon to be a reality? Expert Review of Gastroenterology & Hepatology 2017, 11, 785–788. doi:10.1080/17474124.2017.1342534.

2. Abedon, S.T. Use of phage therapy to treat long-standing, persistent, or chronic bacterial infections. Advanced Drug Delivery Reviews 2018, (in press). doi:10.1016/j.addr.2018.06.018.

3. Schmidt, C. Phage therapy’s latest makeover. Nature Biotechnology 2019, 37, 581–586. doi:10.1038/s41587-019-0133-z.

4. Kortright, K.E.; Chan, B.K.; Koff, J.L.; Turner, P.E. Phage therapy: A renewed approach to combat antibiotic-resistant bacteria. Cell Host & Microbe 2019, 25, 219–232. doi:10.1016/j.chom.2019.01.014.

5. Schooley, R.T.; Biswas, B.; Gill, J.J.; Hernandez-Morales, A.; Lancaster, J.; Lessor, L.; Barr, J.J.; Reed, S.L.; Rohwer, F.; Benler, S.; Segall, A.M.; Taplitz, R.; Smith, D.M.; Kerr, K.; Kumaraswamy, M.; Nizet, V.; Lin, L.; McCauley, M.D.; Strathdee, S.A.; Benson, C.A.; Pope, R.K.; Leroux, B.M.; Picel, A.C.; Mateczun, A.J.; Cilwa, K.E.; Regeimbal, J.M.; Estrella, L.A.; Wolfe, D.M.; Henry, M.S.; Quinones, J.; Salka, S.; Bishop-Lilly, K.A.; Young, R.; Hamilton, T. Development and Use of Personalized Bacteriophage-Based Therapeutic Cocktails To Treat a Patient with a Disseminated Resistant Acinetobacter baumannii Infection. Antimicrobial Agents and Chemotherapy 2017, 61. doi:10.1128/AAC.00954-17.

6. Schooley, R.T.; Biswas, B.; Gill, J.J.; Hernandez-Morales, A.; Lancaster, J.; Lessor, L.; Barr, J.J.; Reed, S.L.; Rohwer, F.; Benler, S.; Segall, A.M.; Taplitz, R.; Smith, D.M.; Kerr, K.; Kumaraswamy, M.; Nizet, V.; Lin, L.; McCauley, M.D.; Strathdee, S.A.; Benson, C.A.; Pope, R.K.; Leroux, B.M.; Picel, A.C.; Mateczun, A.J.; Cilwa, K.E.; Regeimbal, J.M.; Estrella, L.A.; Wolfe, D.M.; Henry, M.S.; Quinones, J.; Salka, S.; Bishop-Lilly, K.A.; Young, R.; Hamilton, T. Erratum for Schooley et al., “Development and Use of Personalized Bacteriophage-Based Therapeutic Cocktails To Treat a Patient with a Disseminated Resistant Acinetobacter baumannii Infection”. Antimicrobial Agents and Chemotherapy 2018, 62. doi:10.1128/AAC.02221-18.

7. Dedrick, R.M.; Guerrero-Bustamante, C.A.; Garlena, R.A.; Russell, D.A.; Ford, K.; Harris, K.; Gilmour, K.C.; Soothill, J.; Jacobs-Sera, D.; Schooley, R.T.; Hatfull, G.F.; Spencer, H. Engineered bacteriophages for treatment of a patient with a disseminated drug-resistant Mycobacterium abscessus. Nature Medicine 2019, 25, 730–733. doi:10.1038/s41591-019-0437-z.

8. Chan, B.K.; Turner, P.E.; Kim, S.; Mojibian, H.R.; Elefteriades, J.A.; Narayan, D. Phage treatment of an aortic graft infected with Pseudomonas aeruginosa. Evolution, Medicine, and Public Health 2018, 2018, 60–66. doi:10.1093/emph/eoy005.

9. Rhoads, D.D.; Wolcott, R.D.; Kuskowski, M.A.; Wolcott, B.M.; Ward, L.S.; Sulakvelidze, A. Bacteriophage therapy of venous leg ulcers in humans: results of a phase I safety trial. Journal of Wound Care 2009, 18, 237–238, 240–243. doi:10.12968/jowc.2009.18.6.42801.

10. Sarker, S.A.; Sultana, S.; Reuteler, G.; Moine, D.; Descombes, P.; Charton, F.; Bourdin, G.; McCallin, S.; Ngom-Bru, C.; Neville, T.; Akter, M.; Huq, S.; Qadri, F.; Talukdar, K.; Kassam, M.; Delley, M.; Loiseau, C.; Deng, Y.; El Aidy, S.; Berger, B.; Brüssow, H. Oral Phage Therapy of Acute Bacterial Diarrhea With Two Coliphage Preparations: A Randomized Trial in Children From Bangladesh. EBioMedicine 2016, 4, 124–137. doi:10.1016/j.ebiom.2015.12.023.

11. Jault, P.; Leclerc, T.; Jennes, S.; Pirnay, J.P.; Que, Y.A.; Resch, G.; Rousseau, A.F.; Ravat, F.; Carsin, H.; Le Floch, R.; Schaal, J.V.; Soler, C.; Fevre, C.; Arnaud, I.; Bretaudeau, L.; Gabard, J. Efficacy and tolerability of a cocktail of bacteriophages to treat burn wounds infected by Pseudomonas aeruginosa (PhagoBurn): a randomised, controlled, double-blind phase 1/2 trial. The Lancet. Infectious Diseases 2019, 19, 35–45. doi:10.1016/S1473-3099(18)30482-1.

12. Smith, H.W.; Huggins, M.B. Successful treatment of experimental *Escherichia coli* infections in mice using phage: its general superiority over antibiotics. Journal of general microbiology 1982, 128, 307–318.

13. Henry, M.; Lavigne, R.; Debarbieux, L. Predicting in vivo efficacy of therapeutic bacteriophages used to treat pulmonary infections. Antimicrobial Agents and Chemotherapy 2013, 57, 5961–5968. doi:10.1128/AAC.01596-13.

14. Arnold, F.H. Innovation by Evolution: Bringing New Chemistry to Life (Nobel Lecture). Angewandte Chemie (International Ed. in English) 2019. doi:10.1002/anie.201907729.

15. D’Herelle, F. Immunity in Natural Infectious Disease, authorized english edition (g h smith) ed.; Williams & Wilkins Co.: Baltimore, MD, USA, 1924.

16. McEwen, S.; Collignon, P. Antimicrobial Resistance: a One Health Perspective. Microbiol Spectrum 2018, 6, ARBA-0009–2017. doi:doi:10.1128/microbiolspec.ARBA-0009-2017.

17. Bull, J.J.; Levin, B.R.; DeRouin, T.; Walker, N.; Bloch, C.A. Dynamics of success and failure in phage and antibiotic therapy in experimental infections. BMC microbiology 2002, 2, 35.

18. Bull, J.J.; Vegge, C.S.; Schmerer, M.; Chaudhry, W.N.; Levin, B.R. Phenotypic resistance and the dynamics of bacterial escape from phage control. PloS One 2014, 9, e94690. doi:10.1371/journal.pone.0094690.

19. Merril, C.R.; Biswas, B.; Carlton, R.; Jensen, N.C.; Creed, G.J.; Zullo, S.; Adhya, S. Long-circulating bacteriophage as antibacterial agents. Proceedings of the National Academy of Sciences of the United States of America 1996, 93, 3188–3192.

20. Sutherland, I.W.; Hughes, K.A.; Skillman, L.C.; Tait, K. The interaction of phage and biofilms. FEMS microbiology letters 2004, 232, 1–6.

21. Lu, T.K.; Collins, J.J. Dispersing biofilms with engineered enzymatic bacteriophage. Proceedings of the National Academy of Sciences of the United States of America 2007, 104, 11197–11202. doi:10.1073/pnas.0704624104.

22. Chan, B.K.; Abedon, S.T. Phage therapy pharmacology: phage cocktails. In Advances in Applied Microbiology; Laskin, A.I.; Sariaslani, S.; Gadd, G.M., Eds.; Academic Press, 2012; Vol. 78, pp. 1–23. doi:10.1016/B978-0-12-394805-2.00001-4.

23. Adams, M.H. Bacteriophages; Interscience Publishers: New York, NY, 1959.

24. Campbell, A. Conditions for the existence of bacteriophage. Evolution 1961, 15, 143–165.

25. Levin, B.R.; Stewart, F.M.; Chao, L. Resource - limited growth, competition, and predation: a model and experimental studies with bacteria and bacteriophage. The American Naturalist 1977, 977, 3–24.

26. Roach, D.R.; Leung, C.Y.; Henry, M.; Morello, E.; Singh, D.; Di Santo, J.P.; Weitz, J.S.; Debarbieux, L. Synergy between the Host Immune System and Bacteriophage Is Essential for Successful Phage Therapy against an Acute Respiratory Pathogen. Cell Host & Microbe 2017, 22, 38–47.e4. doi:10.1016/j.chom.2017.06.018.

27. Abedon, S.T. Bacteriophage-mediated biocontrol of wound infections, and ecological exploitation of biofilms by phages. In Recent Clinical Techniques, Results, and Research in Wounds; Springer International Publishing AG, 2018; pp. 1–38.

28. Darch, S.E.; Kragh, K.N.; Abbott, E.A.; Bjarnsholt, T.; Bull, J.J.; Whiteley, M. Phage inhibit pathogen dissemination by targeting bacterial migrants in a chronic infection model. mBio 2017, 8. doi:10.1128/mBio.00240-17.

29. Bull, J.J. Optimality models of phage life history and parallels in disease evolution. Journal of Theoretical Biology 2006, 241, 928–938. doi:10.1016/j.jtbi.2006.01.027.

30. Heineman, R.H.; Springman, R.; Bull, J.J. Optimal foraging by bacteriophages through host avoidance. The American Naturalist 2008, 171, E149–157. doi:10.1086/528962.

31. Westwater, C.; Kasman, L.M.; Schofield, D.A.; Werner, P.A.; Dolan, J.W.; Schmidt, M.G.; Norris, J.S. Use of genetically engineered phage to deliver antimicrobial agents to bacteria: an alternative therapy for treatment of bacterial infections. Antimicrobial agents and chemotherapy 2003, 47, 1301–1307.

32. Bull, J.J.; Regoes, R.R. Pharmacodynamics of non-replicating viruses, bacteriocins and lysins. Proceedings. Biological Sciences / The Royal Society 2006, 273, 2703–2712. doi:10.1098/rspb.2006.3640.

33. Hodyra-Stefaniak, K.; Lahutta, K.; Majewska, J.; Kaźmierczak, Z.; Lecion, D.; Harhala, M.; Kęska, W.; Owczarek, B.; Jończyk-Matysiak, E.; Kłopot, A.; Miernikiewicz, P.; Kula, D.; Górski, A.; Dąbrowska, K. Bacteriophages engineered to display foreign peptides may become short-circulating phages. Microbial Biotechnology 2019, 12, 730–741. doi:10.1111/1751-7915.13414.

34. Sutherland, I.W.; Wilkinson, J.F. Depolymerases for bacterial exopolysaccharides obtained from phage-infected bacteria. Journal of General Microbiology 1965, 39, 373–383.

35. Sutherland, I.W. Polysaccharide lyases. FEMS microbiology reviews 1995, 16, 323–347.

36. Hughes, K.A.; Sutherland, I.W.; Clark, J.; Jones, M.V. Bacteriophage and associated polysaccharide depolymerases–novel tools for study of bacterial biofilms. Journal of Applied Microbiology 1998, 85, 583–590.

37. Hanlon, G.W.; Denyer, S.P.; Olliff, C.J.; Ibrahim, L.J. Reduction in exopolysaccharide viscosity as an aid to bacteriophage penetration through Pseudomonas aeruginosa biofilms. Applied and environmental microbiology 2001, 67, 2746–2753. doi:10.1128/AEM.67.6.2746-2753.2001.

38. Azeredo, J.; Sutherland, I.W. The use of phages for the removal of infectious biofilms. Current Pharmaceutical Biotechnology 2008, 9, 261–266.

39. Bull, J.J.; Vimr, E.R.; Molineux, I.J. A tale of tails: Sialidase is key to success in a model of phage therapy against K1-capsulated Escherichia coli. Virology 2010, 398, 79–86.

40. Mushtaq, N.; Redpath, M.B.; Luzio, J.P.; Taylor, P.W. Prevention and cure of systemic Escherichia coli K1 infection by modification of the bacterial phenotype. Antimicrobial agents and chemotherapy 2004, 48, 1503–1508.

41. Mushtaq, N.; Redpath, M.B.; Luzio, J.P.; Taylor, P.W. Treatment of experimental Escherichia coli infection with recombinant bacteriophage-derived capsule depolymerase. The Journal of Antimicrobial Chemotherapy 2005, 56, 160–165. doi:10.1093/jac/dki177.

42. Lin, T.L.; Hsieh, P.F.; Huang, Y.T.; Lee, W.C.; Tsai, Y.T.; Su, P.A.; Pan, Y.J.; Hsu, C.R.; Wu, M.C.; Wang, J.T. Isolation of a bacteriophage and its depolymerase specific for K1 capsule of Klebsiella pneumoniae: implication in typing and treatment. The Journal of Infectious Diseases 2014, 210, 1734–1744. doi:10.1093/infdis/jiu332.

43. Lin, H.; Paff, M.L.; Molineux, I.J.; Bull, J.J. Therapeutic application of phage capsule depolymerases against K1, K5, and K30 capsulated E. coli in mice. Frontiers in Microbiology 2017, 8, 2257. doi:10.3389/fmicb.2017.02257.

44. Lin, H.; Paff, M.L.; Molineux, I.J.; Bull, J.J. Antibiotic therapy using phage depolymerases: Robustness across a range of conditions. Viruses 2018, 10. doi:10.3390/v10110622.

45. Cornelissen, A.; Ceyssens, P.J.; T’Syen, J.; Van Praet, H.; Noben, J.P.; Shaburova, O.V.; Krylov, V.N.; Volckaert, G.; Lavigne, R. The T7-related Pseudomonas putida phage *Ω*15 displays virion-associated biofilm degradation properties. PloS One 2011, 6, e18597. doi:10.1371/journal.pone.0018597.

46. Donlan, R.M.; Costerton, J.W. Biofilms: survival mechanisms of clinically relevant microorganisms. Clinical microbiology reviews 2002, 15, 167–193.

47. Tseng, B.S.; Zhang, W.; Harrison, J.J.; Quach, T.P.; Song, J.L.; Penterman, J.; Singh, P.K.; Chopp, D.L.; Packman, A.I.; Parsek, M.R. The extracellular matrix protects Pseudomonas aeruginosa biofilms by limiting the penetration of tobramycin. Environmental Microbiology 2013, 15, 2865–2878. doi:10.1111/1462-2920.12155.

48. Pei, R.; Lamas-Samanamud, G.R. Inhibition of biofilm formation by T7 bacteriophages producing quorum-quenching enzymes. Applied and Environmental Microbiology 2014, 80, 5340–5348. doi:10.1128/AEM.01434-14.

49. Bárdy, P.; Pantuůček, R.; Benešíkk, M.; Doškarř, J. Genetically modified bacteriophages in applied microbiology. Journal of Applied Microbiology 2016, 121, 618–633. doi:10.1111/jam.13207.

50. Bessler, W.; Fehmel, F.; Freund-Mölbert, E.; Knüfermann, H.; Stirm, S. Escherichia coli capsule bacteriophages. IV. Free capsule depolymerase 29. Journal of Virology 1975, 15, 976–984.

51. Kassa, T.; Chhibber, S. Thermal treatment of the bacteriophage lysate of Klebsiella pneumoniae B5055 as a step for the purification of capsular depolymerase enzyme. Journal of Virological Methods 2012, 179, 135–141. doi:10.1016/j.jviromet.2011.10.011.

52. Schmerer, M.; Molineux, I.J.; Bull, J.J. Synergy as a rationale for phage therapy using phage cocktails. PeerJ 2014, 2, e590. doi:10.7717/peerj.590.

53. Gladstone, E.G.; Molineux, I.J.; Bull, J.J. Evolutionary principles and synthetic biology: avoiding a molecular tragedy of the commons with an engineered phage. Journal of Biological Engineering 2012, 6, 13. doi:10.1186/1754-1611-6-13.

54. Hashemolhosseini, S.; Holmes, Z.; Mutschler, B.; Henning, U. Alterations of receptor specificities of coliphages of the T2 family. Journal of Molecular Biology 1994, 240, 105–110.

55. Hashemolhosseini, S.; Holmes, Z.; Mutschler, B.; Henning, U. Alterations of receptor specificities of coliphages of the T2 family. Journal of Molecular Biology 1994, 240, 105–110. doi:10.1006/jmbi.1994.1424.

56. Dinsmore, P.K.; Klaenhammer, T.R. Bacteriophage resistance in Lactococcus. Molecular Biotechnology 1995, 4, 297–314. doi:10.1007/BF02779022.

57. Durmaz, E.; Klaenhammer, T.R. Abortive phage resistance mechanism AbiZ speeds the lysis clock to cause premature lysis of phage-infected Lactococcus lactis. Journal of Bacteriology 2007, 189, 1417–1425. doi:10.1128/JB.00904-06.

58. Labrie, S.J.; Samson, J.E.; Moineau, S. Bacteriophage resistance mechanisms. Nature Reviews. Microbiology 2010, 8, 317–327. doi:10.1038/nrmicro2315.

59. Levin, B.R. Frequency-dependent selection in bacterial populations. Philosophical Transactions of the Royal Society of London. Series B, Biological Sciences 1988, 319, 459–472. doi:10.1098/rstb.1988.0059.

60. Bohannan, B.J.M.; Lenski, R.E. Linking genetic change to community evolution: insights from studies of bacteria and bacteriophage. Ecology Letters 2000, 3, 362–377. doi:10.1046/j.1461-0248.2000.00161.x.

61. Mizoguchi, K.; Morita, M.; Fischer, C.R.; Yoichi, M.; Tanji, Y.; Unno, H. Coevolution of bacteriophage PP01 and Escherichia coli O157:H7 in continuous culture. Applied and Environmental Microbiology 2003, 69, 170–176. doi:10.1128/aem.69.1.170-176.2003.

62. Gómez, P.; Buckling, A. Bacteria-phage antagonistic coevolution in soil. Science (New York, N.Y.) 2011, 332, 106–109. doi:10.1126/science.1198767.

63. Díaz-Muñoz, S.L.; Koskella, B. Bacteria-phage interactions in natural environments. Advances in Applied Microbiology 2014, 89, 135–183. doi:10.1016/B978-0-12-800259-9.00004-4.

64. Fortuna, M.A.; Barbour, M.A.; Zaman, L.; Hall, A.R.; Buckling, A.; Bascompte, J. Coevolutionary dynamics shape the structure of bacteria-phage infection networks. Evolution; International Journal of Organic Evolution 2019, 73, 1001–1011. doi:10.1111/evo.13731.

65. Harcombe, W.R.; Bull, J.J. Impact of phages on two-species bacterial communities. Applied and Environmental Microbiology 2005, 71, 5254–5259. doi:10.1128/AEM.71.9.5254-5259.2005.

66. Hernandez, C.A.; Koskella, B. Phage resistance evolution in vitro is not reflective of in vivo outcome in a plant-bacteria-phage system. Evolution; International Journal of Organic Evolution 2019. doi:10.1111/evo.13833.

67. LaVergne, S.; Hamilton, T.; Biswas, B.; Kumaraswamy, M.; Schooley, R.T.; Wooten, D. Phage therapy for a multidrug-resistant *Acinetobacter baumannii* craniectomy site Infection. Open Forum Infectious Diseases 2018, 5, ofy064. doi:10.1093/ofid/ofy064.

68. Yoichi, M.; Abe, M.; Miyanaga, K.; Unno, H.; Tanji, Y. Alteration of tail fiber protein gp38 enables T2 phage to infect Escherichia coli O157:H7. Journal of Biotechnology 2005, 115, 101–107. doi:10.1016/j.jbiotec.2004.08.003.

69. Mahichi, F.; Synnott, A.J.; Yamamichi, K.; Osada, T.; Tanji, Y. Site-specific recombination of T2 phage using IP008 long tail fiber genes provides a targeted method for expanding host range while retaining lytic activity. FEMS microbiology letters 2009, 295, 211–217. doi:10.1111/j.1574-6968.2009.01588.x.

70. Pouillot, F.; Blois, H.; Iris, F. Genetically engineered virulent phage banks in the detection and control of emergent pathogenic bacteria. Biosecurity and Bioterrorism: Biodefense Strategy, Practice, and Science 2010, 8, 155–169. doi:10.1089/bsp.2009.0057.

71. Korona, R.; Levin, B.R. Phage-mediated selection and the evolution and maintenance of restriction-modification. Evolution; International Journal of Organic Evolution 1993, 47, 556–575. doi:10.1111/j.1558-5646.1993.tb02113.x.

72. Chan, B.K.; Sistrom, M.; Wertz, J.E.; Kortright, K.E.; Narayan, D.; Turner, P.E. Phage selection restores antibiotic sensitivity in MDR Pseudomonas aeruginosa. Scientific Reports 2016, 6, 26717. doi:10.1038/srep26717.

73. Gurney, J.; Pleška, M.; Levin, B.R. Why put up with immunity when there is resistance: an excursion into the population and evolutionary dynamics of restriction-modification and CRISPR-Cas. Philosophical Transactions of the Royal Society of London. Series B, Biological Sciences 2019, 374, 20180096. doi:10.1098/rstb.2018.0096.

74. Chaudhry, W.N.; Concepción-Acevedo, J.; Park, T.; Andleeb, S.; Bull, J.J.; Levin, B.R. Synergy and order effects of antibiotics and phages in killing Pseudomonas aeruginosa biofilms. PloS One 2017, 12, e0168615. doi:10.1371/journal.pone.0168615.

75. Uchiyama, J.; Shigehisa, R.; Nasukawa, T.; Mizukami, K.; Takemura-Uchiyama, I.; Ujihara, T.; Murakami, H.; Imanishi, I.; Nishifuji, K.; Sakaguchi, M.; Matsuzaki, S. Piperacillin and ceftazidime produce the strongest synergistic phage-antibiotic effect in Pseudomonas aeruginosa. Archives of Virology 2018, 163, 1941–1948. doi:10.1007/s00705-018-3811-0.

76. Segall, A.M.; Roach, D.R.; Strathdee, S.A. Stronger together? Perspectives on phage-antibiotic synergy in clinical applications of phage therapy. Current Opinion in Microbiology 2019, 51, 46–50. doi:10.1016/j.mib.2019.03.005.

77. Abedon, S.T. Phage-antibiotic combination treatments: Antagonistic impacts of antibiotics on the pharmacodynamics of phage therapy? Antibiotics (Basel, Switzerland) 2019, 8, (in press).

78. Matsuda, T.; Freeman, T.A.; Hilbert, D.W.; Duff, M.; Fuortes, M.; Stapleton, P.P.; Daly, J.M. Lysis-deficient bacteriophage therapy decreases endotoxin and inflammatory mediator release and improves survival in a murine peritonitis model. Surgery 2005, 137, 639–646. doi:10.1016/j.surg.2005.02.012.

